# Lysine acetylation regulates antibiotic resistance in *Escherichia coli*

**DOI:** 10.1101/2022.02.22.481468

**Authors:** Zuye Fang, Fubin Lai, Kun Cao, Ziyuan Zhang, Linlin Cao, Shiqin Liu, Yufeng Duan, Xingfeng Yin, Ruiguang Ge, Qing-Yu He, Xuesong Sun

## Abstract

Antibiotic resistance is increasingly becoming a serious challenge to public health. The regulation of metabolism by post-translational modifications (PTMs) has been widely studied; however, the comprehensive mechanism underlying the regulation of acetylation in bacterial resistance against antibiotics is unknown. Herein, with *Escherichia coli* as the model, we performed quantitative analysis of the acetylated proteome of wild-type sensitive strain (WT) and ampicillin- (Re-Amp), kanamycin- (Re-Kan), and polymyxin B-resistant (Re-Pol) strains. Based on bioinformatics analysis combined with biochemical validations, we found that a common regulatory mechanism exists between the different resistant strains. Acetylation negatively regulates bacterial metabolism to maintain antibiotic resistance, but positively regulates bacterial motility. Further analyses revealed that key enzymes in various metabolic pathways were differentially acetylated. Particularly, pyruvate kinase (PykF), a key glycolytic enzyme regulating bacterial metabolism, and its acetylated form were highly expressed in the three resistant types and were identified as reversibly acetylated by the deacetylase CobB and the acetyl-transferase PatZ, and also could be acetylated by non-enzyme AcP in vitro. Further, the deacetylation of Lys413 of PykF increased the enzyme activity by changing the conformation of ATP binding site of PykF, resulting in an increase in energy production, which in turn increased the sensitivity of drug-resistant strains to antibiotics. This study provides novel insights for understanding bacterial resistance and lays the foundation for future research on regulation of acetylation in antibiotic-resistant strains.

**Importance:** The misuse of antibiotics has resulted in an emergence of a large number of antibiotic-resistant strains, which seriously threaten human health. Bacterial metabolism is tightly controlled by protein post-translational modifications, especially acetylation. However, the comprehensive mechanism underlying regulation of acetylation in bacterial resistance remains unexplored. Here, acetylation was found to positively regulate bacterial motility and negatively regulate energy metabolism, which was common in all the different antibiotic-resistant strains. Moreover, the acetylation and deacetylation process of PykF was uncovered, and deacetylation of the Lys 413 of PykF was found to contribute to bacterial sensitivity to antibiotics. This study provides a new direction for research on development of bacterial resistance through post-translational modifications and provides a theoretical basis for the development of antibacterial drugs.

## INTRODUCTION

Antibiotics are being widely used to treat infections and have significantly reduced the incidences of infection, improving the quality of life and reducing mortality. Unfortunately, the frequent abuse of antibiotics has led to a rapid emergence of antibiotic-resistant strains, which are a serious threat to human health(Baron et al., 2014). Bacteria have developed different mechanisms to resist different antibiotics. For example, the production of highly active penicillin binding protein to resist ampicillin(Li et al., 2019), structural change in the 30S subunit to resist kanamycin (Alangaden et al., 1998; Kang et al., 2012), and structural alteration of lipid A to combat with polymyxin B (Olaitan et al., 2014; Yu et al., 2015).

Protein post-translational modifications (PTMs) regulate many important biological pathways in bacteria. In recent years, with the development of mass spectrometry (MS) and enrichment strategies, numerous PTMs have been found to be related to bacterial metabolism(Colak et al., 2013; Hentchel & Escalante-Semerena, 2015; Simithy et al., 2017; Sreedhar et al., 2020; Yang et al., 2015; J. Zhang et al., 2009). Especially, with the ongoing research on acetylation, many key kinases have been found to be regulated by acetylation. The extent of acetylation in *E. coli* varies at different growth stages, and is higher in the stationary phase than that in the logarithmic growth phase, suggesting that the change in acetylation affects the bacterial metabolic state(Weinert et al., 2017). Lysine acetylation is one of the most widely known forms of PTMs of proteins in bacteria, which was involved in the regulation of bacterial transcription, translation, metabolism, stress response, and other important physiological processes (Hentchel & Escalante-Semerena, 2015; Ma & Wood, 2011; Ren et al., 2017). In *E. coli*, two mechanisms of Nε-lysine acetylation are known. The main mechanism is acetyl phosphate (AcP) dependent nonenzymatic acetylation. AcP is an intermediate product of the phosphotransferase (Pta)-acetate kinase (AckA) pathway, which can mutually convert acetyl-CoA (AcCoA), inorganic phosphate and ADP into acetate, CoA and ATP. The secondary mechanism is enzymatic reaction mediated by Nε-lysine acetyltransferase (KAT) and deacetylases (KDACs) (Weinert et al., 2013). The most common KAT in bacteria is PatZ, also called as Pka and YfiQ(Kuhn et al., 2014b; Starai & Escalante-Semerena, 2004; Weinert et al., 2013). In addition, four new types of KATs, RimI, YiaC, YjaB, and PhnO, ware also discovered(Christensen et al., 2018). NAD^+^ depedennt CobB is considered as the common KDAC in bacteria. When *cobB* of *E. coli* was deleted, acetylation levels were increased and resistance ability to heat shock and oxidative stress was improved (Ma & Wood, 2011; Weinert et al., 2013). The same phenomenon has also been observed in mycobacteria(Liu et al., 2014).

In bacteria, the activity of multiple transcription factors including RcsB, Lrp and PhoP can be regulated by acetylation (Ren et al., 2017; Ren et al., 2016). A large number of acetylated proteins are also involved in the important carbon metabolism pathways of bacteria such as tricarboxylic acid (TCA) cycle, glycolysis, and oxidative phosphorylation (Hentchel & Escalante-Semerena, 2015; Ma & Wood, 2011; Smith et al., 2018). This shows that acetylation will significantly influence the regulation of bacterial metabolism. In addition, acetylation also plays an important role in the regulation of bacterial pathogenicity and virulence (Ren et al., 2017). For example, in *Salmonella*, PhoP can be acetylated by Pat at lysine 201 (K201) and the deacetylation of PhoP K201 can ensure that the bacteria adapt to the oxidative stress derived from the host cells. This study indicates that the acetylation at this site is necessary for the pathogenicity of the bacteria(Ren et al., 2016). Although regulation of bacterial metabolism by PTMs has been reported before, how lysine acetylation regulates the bacterial resistance to antibiotics has not yet been deciphered.

This study aims to reveal common comprehensive mechanism to regulate antibiotic resistance of bacteria against different types of antibiotics. Here, we used quantitative proteomics analysis combined with antibody enrichment to investigate how acetylation affects bacterial resistance to different types of antibiotics including β-lactams (ampicillin), aminoglycosides (kanamycin), and polypeptides (polymyxin B). Further, biochemical analyses were used to unravel the key biological functions of the specific acetylated site/s in proteins involved to reveal novel comprehensive mechanisms of PTMs in regulating bacterial resistance. This study could pave the way for the development of new drugs to treat infections caused by the drug-resistant strains.

## RESULTS

### Acetylation intensities in different antibiotic-resistant *Escherichia coli*

The differences in the acetylation intensities of the whole proteome between the wild type (WT) and *Escherichia coli* resistant to different antibiotics were investigated. To generate domesticated highly resistant strains, *E. coli* BW25113 was continuously subcultured in media with ampicillin, kanamycin, and polymyxin B, the representative drugs of the antibiotic classes β-lactams, aminoglycosides, and polypeptides, respectively. The measurement of the minimum inhibitory concentration (MIC) showed that the MIC of the highly resistant bacterial strains was increased approximately 60-fold when compared with that of the WT. For ampicillin, the MICs of WT and Re-Amp were 8.2 μg/mL and 481.7 μg/ mL, respectively; for kanamycin, the MICs of WT and Re-Kan were 0.5 μg/mL and 31.3 μg/mL, respectively, and for polymyxin B, the MICs of WT and Re-Pol were 0.3 μg/mL and 22.6 μg/mL, respectively (Figure 1A).

**Figure 1.**
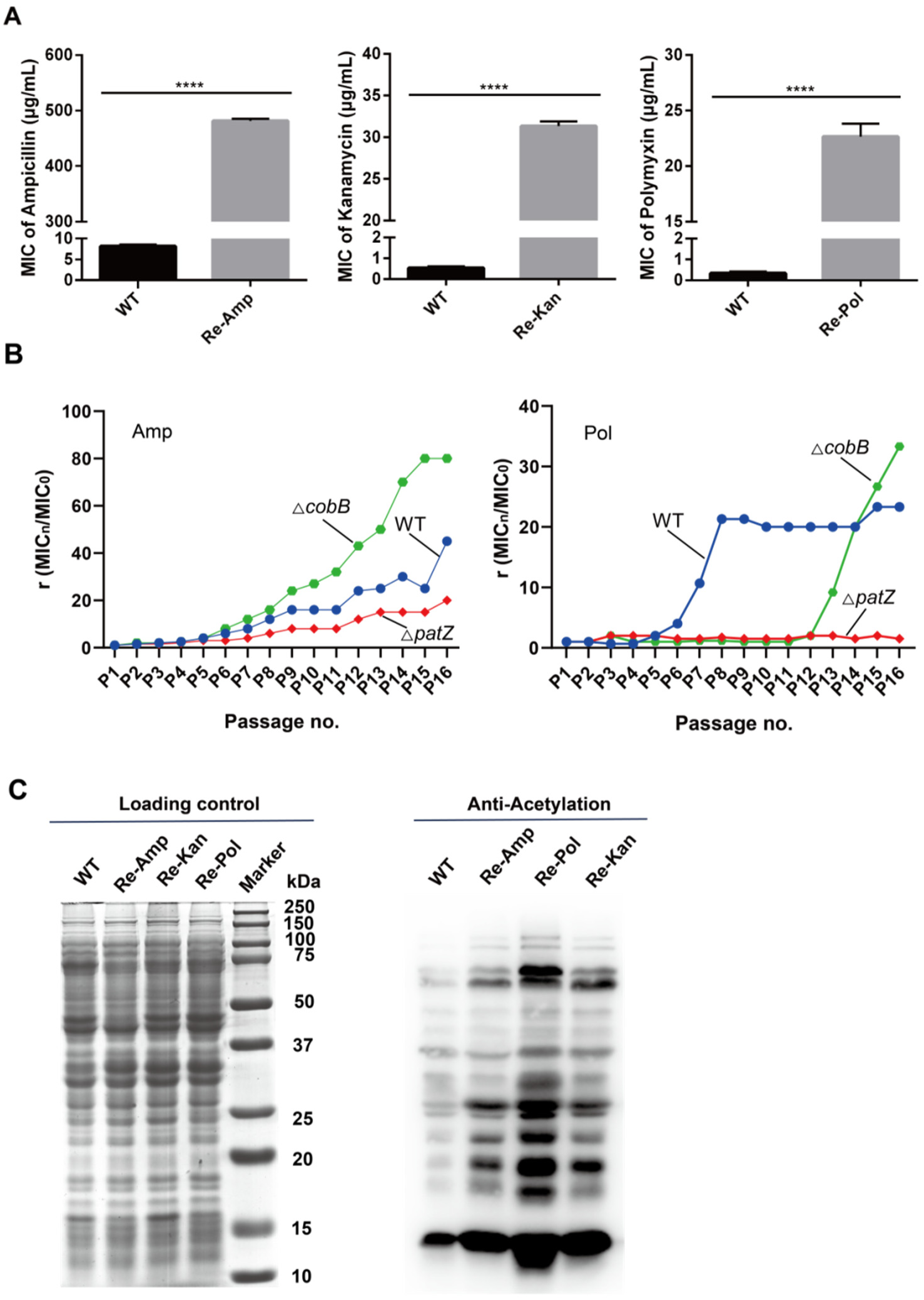
The relationship between protein acetylation modification of *Escherichia coli* and the formation of drug resistance. (A) The average MIC of the highly resistant strains from left to right in the figure are: for Amp (WT=8.2 μg/mL, Re-Amp=481.7 μg/mL), for Kan (WT=0.5 μg/mL, Re-Kan=31.3 μg/mL), and for Pol (WT=0.3 μg/mL, Re-Pol=22.6 μg/mL) (n = 3 independent biological replicates). Two-tailed unpaired t-test was used for statistical analysis. (*, p < 0.05; **, p < 0.01; ***, p < 0.005; ****, p < 0.001). (B) The development of resistance formation of *Escherichia coli* BW25113 and Δ*patZ* and Δ*cobB* strains to ampicillin (left) and polymyxin (right). (C) Immunoblot of the *Escherichia coli* lysate with the α-AcK monoclonal antibody. Figure 1—source data 1. MIC and MIC fold change for Figure 1A-B Figure 1—source data 2. Uncropped gels and blots for Figure 1C.

We collected WT and the three antibiotic-resistant strains in a stable growth period (OD_600_ ∼ 0.8) in LB medium as the accumulation of acetyl phosphate, the acetylation level at this stage is higher than that in other stages (Weinert et al., 2017). Western blotting with α-AcK monoclonal antibody indicated a certain difference in the acetylation intensities of the whole protein level between the WT and the three antibiotic-resistant strains (Figure 1C). In particular, the proteins of drug-resistant bacteria showed a higher level of acetylation than that of WT, suggesting that protein acetylation may be conducive to the maintenance of drug resistance. In addition, different drug-resistant bacteria showed different acetylated protein profiles, indicating that different kinds of drug-resistant bacteria may also have the specific regulation mechanism of drug resistance. In order to further explore the role of acetylation in antibiotic resistance, Δ*patZ* and Δ*cobB* were used to study the developing ability of resistance to ampicillin and polymyxin B (Figure 1B). The results showed that Δ*cobB* was more likely to form antibiotic resistance while Δ*patZ* was on the contrary. Interestingly, the resistance to polymyxin B of Δ*cobB* was latent and increased suddenly, which may be related to the complicate action mechanism of polymyxin B (Trimble et al., 2016). This result indicated that acetylation may positively correlate with antibiotic resistance of bacteria.

### Distribution of acetylation in *Escherichia coli* is stable

To further study the relationship between acetylation and antibiotic resistance, acetylated peptides of the WT and the three antibiotic-resistant strains were enriched with α-AcK antibody resin and identified with DIA-based LC-MS/MS. All the distribution of mass error for precursor ions was <0.03 Da, indicating acceptable mass accuracy of the MS data. From three independent biological replicates, 2971 lysine acetylated sites of 1039 proteins, 3108 acetylated sites of 1035 proteins, 3282 acetylated sites of 1085 proteins, and 3689 lysine acetylated sites of 1115 proteins were identified in WT, Re-Amp, Re-Kan, and Re-Pol, respectively. Hierarchical cluster analysis showed that the proteomic data had good biological reproducibility (Figure 2D-F). Compared with a previous acetyl-proteomic survey in the same species *E. coli* BW25113 which identified 809 acetylated proteins with 2803 modification sites (Castano-Cerezo et al., 2014), our study showed higher enrichment efficiency and MS sensitivity. In total, we identified a total of 1228 acetylated proteins and 4193 acetylated sites in all samples, of which 901 (73.4%) acetylated proteins and 2411 (57.5%) acetylated sites were commonly identified in the sensitive bacteria and three resistant types showing the stability of acetylation in bacteria (Figure 2A). We further assessed the distribution of lysine acetylated sites, and found that most of the proteins were monoacetylated, which was in accord with the acetylome studies in other bacterial species (Liu et al., 2018; Meng et al., 2016; Ouidir et al., 2015). In addition, the distribution of monoacetylated proteins and polyacetylated proteins was almost the same in WT and the resistant strains (Figure 2C). The stable distribution of acetylation in *E. coli* in different resistant phenotypes exhibited the high conservation of this modification.

**Figure 2.**
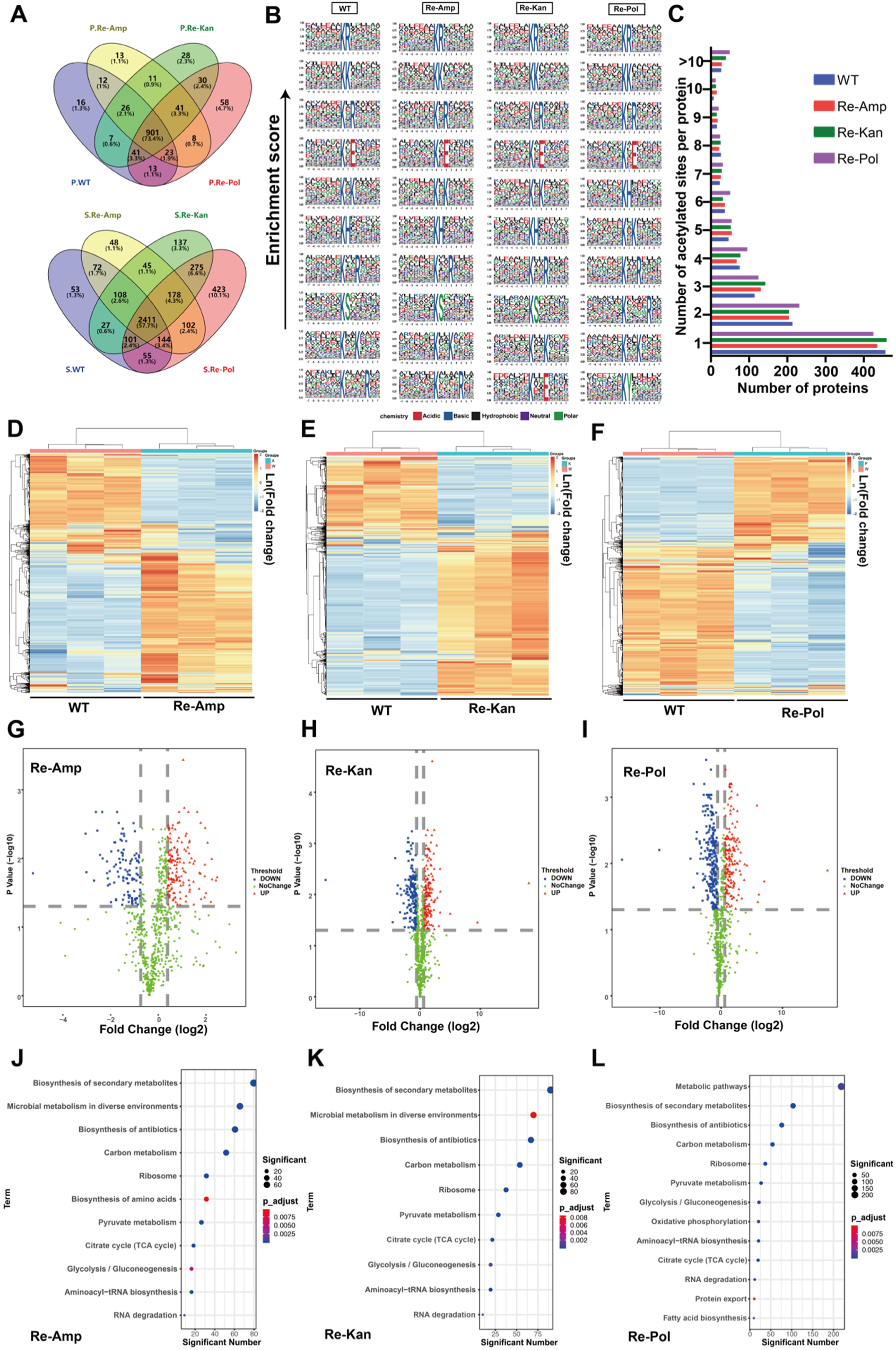
Acetylated proteins and sites in *Escherichia coli*. (A) Statistics of acetylated protein and acetylated sites in the different resistant types. (P, the acetylated protein; S, the acetylated site). (B) Motif significant enrichment analysis on the identified acetylated peptides (7 upstream and downstream amino acids of acetylated lysine). From left to right, WT, Re-Amp, Re-Kan, Re-Pol. P < 0.000001 was considered as significant. (C) The number of acetylated sites identified per protein. (D-F) Heatmap of hierarchical cluster analysis for differential acetylated protein sites. (G–I) Volcano graphs for the statistical analysis of acetylation sites. (the difference multiple is 1.5 times up and down). From left to right, WT, Re-Amp, Re-Kan, Re-Pol. (J–L) KEGG pathway analysis of the differentially acetylated proteins. p < 0.05. From left to right, WT, Re-Amp, Re-Kan, Re-Pol. (A-L) (n = 3 independent biological replicates). Figure 2—source data 1. Quantitative data and KEGG analysis of differential acetylated proteins for Figure 2.

To determine the preference for acetylation of lysine in different antibiotic-resistant strains, the Motif-X tool was used to perform sequence analysis of the acetylated peptides (seven upstream and seven downstream amino acids of the acetylated lysine). Eight motif sequences were overrepresented in all the identified peptides (Figure 2B), including KR/K/H, KxK/R/E, and KxxK/R (K, the acetylated lysine; x, random amino acid residues). The amino acids in these motifs were hydrophilic, indicating that KATs prefer hydrophilic environment and that the occurrence of acetylation sites is stable (Kuhn et al., 2014a). Arginine, lysine, and histidine are preferentially located at the +1 position; lysine, arginine and glutamic acid at the +2 position; and lysine and arginine at the +3 position. Obviously, the +1 position is preferentially a positively charged amino acid, which may be due to that most acetylated proteins were AcP-dependent acetylation. This preference has also been found in other organisms, for example, the KH motif has been identified in humans, plants, and *Vibrio parahaemolyticus* (Fang et al., 2015; Kim et al., 2006; Pan et al., 2014; J. M. Zhang et al., 2009) implying that this kind of protein modification may be relatively conservative. Interestingly, new motifs were detected in this study: KxxE in Re-Kan and KY in Re-Pol, suggesting that bacteria may have new substrate-specific KATs involved in the regulation of resistance owing to the continuous pressure of kanamycin and polymyxin B.

### Maintenance of antibiotic resistance to different antibiotics may depend on a common acetylation regulatory pathway

In order to find out the changes in acetylation of proteins really related to drug resistance of bacteria, the quantitative results of the whole proteome were used as background to subtract. The statistical analysis of the different acetylated peptides (up and down 1.5 folds) in the three resistant strains, showed that, compared with the WT, Re-Amp had 366 differentially acetylated proteins (DAPs), while Re-Kan had 455 DAPs, and Re-Pol had 522 DAPs (Figure 2G-I). Re-Pol possessed more differentially expressed acetylated peptides.

Next, we conducted the KEGG pathway enrichment analysis of the DAPs of the three resistant strains, and found that the DAPs were significantly enriched in the ribosome, aminoacyl-tRNA biosynthesis, RNA degradation, pyruvate metabolism, methane metabolism, glycolysis, TCA cycle, oxidative phosphorylation (Figure 2J-L), which is consistent with the acetylation mainly occurring in the translation and metabolism of prokaryotes and eukaryotes(Hentchel & Escalante-Semerena, 2015). These results further indicate that maintaining bacterial resistance may be related to protein synthesis and energy metabolism regulated by acetylation.

### Acetylation reduces the metabolic activity of *E. coli* to maintain antibiotic resistance

To further investigate how lysine acetylation regulates the bacterial resistance, we conducted a statistical analysis of the identified acetylation sites to find the common sites in the three resistant types. Compared with WT, the acetylation levels of 185 acetylated proteins were simultaneously up-regulated (Co-Up), and 102 acetylated proteins were down-regulated (Co-Down) in all the three resistant strains. KEGG pathway analysis of these commonly changed acetylation sites showed that Co-Down acetylation mainly occurred in ribosome, RNA degradation, signal transduction, and protein export (Figure 3A). Lower levels of acetylation may inhibit signal transduction and prevent antibiotics from binding to the target proteins. Moreover, the acetylation levels of three flagellin proteins associated with bacterial movement were down-regulated, including FliA, FlhD and FliC (Chilcott & Hughes, 2000; Liu & Matsumura, 1996). The decreased bacterial motility is beneficial for bacteria to save energy and resist antibiotic pressure. These results suggest that acetylation may positively regulate bacterial motility. It was observed that the Co-Up of acetylation sites mainly occurs in the TCA cycle, pyruvate metabolism, pentose phosphate pathway, and glycolysis/gluconeogenesis (Figure 3A). Our results further indicated that acetylation may negatively regulate energy metabolism, and an increase in the level of acetylation in resistant strains will lead to a reduced energy metabolism which is beneficial to the development of resistance to the antibiotics.

**Figure 3.**
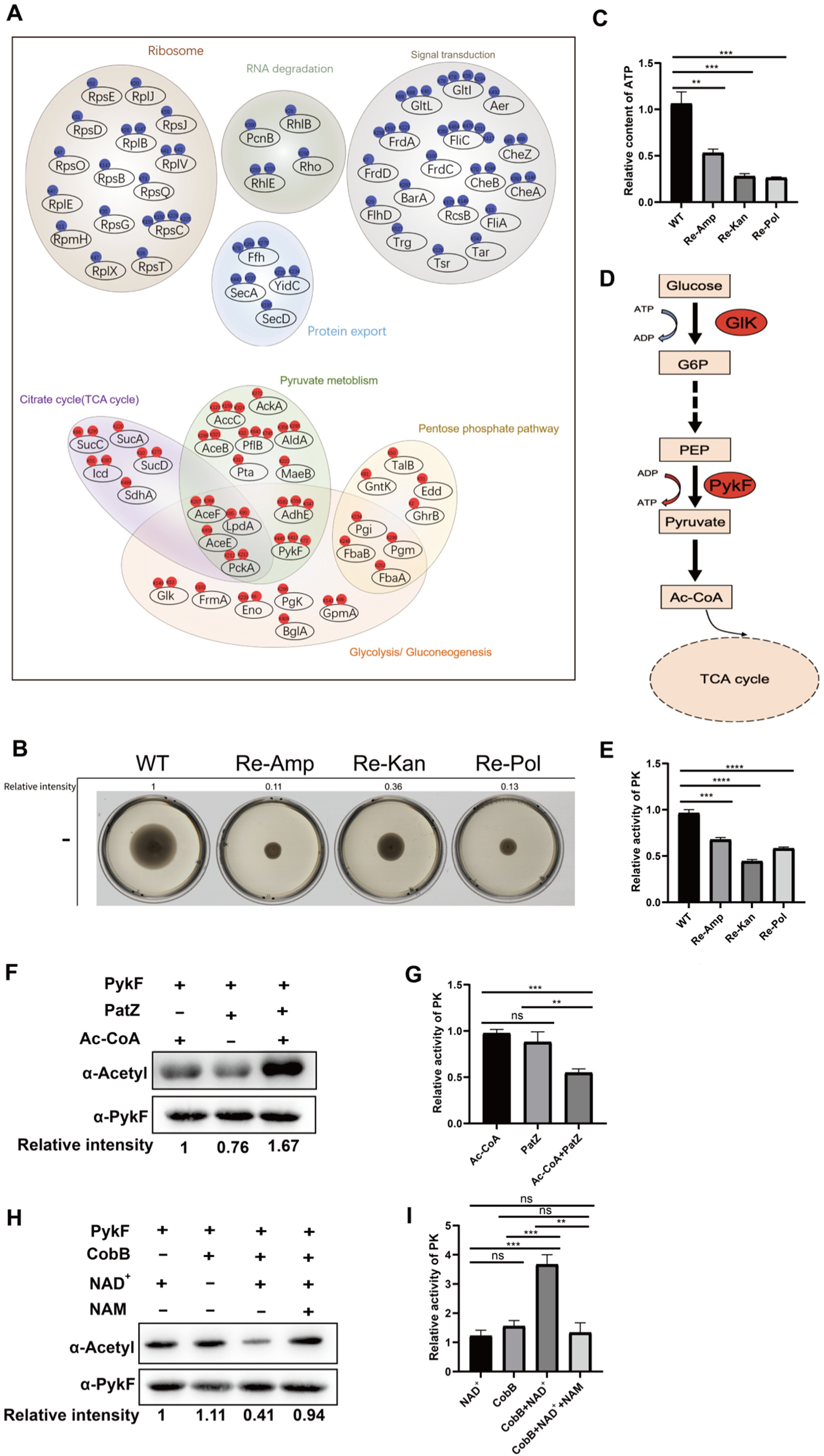
Acetylation negatively regulates the metabolism of antibiotic-resistant strains. (A) KEGG pathway analysis of the Co-Up (red) and Co-Down (blue) acetylated proteins. p < 0.05. (B) Detection of bacterial motility. Relative intensity is the multiple of the bacterial movement area measured by the Image J tool relative to the WT. (C) ATP content detection of resistant strains relative to WT. (D) Key kinases involved in the glycolysis process overexpressed in the three resistant strains. The red oval represents up-regulation in the resistant strains. (E, G, I) PykF enzyme activity detection. Experiments were performed in triplicate. (F) PatZ acetylates PykF in vitro. Western blots are representative of at least three independent replicates. (H) CobB deacetylates PykF in vitro. Western blots are representative of at least three independent replicates. (C, E, G, I) (n = 3 independent biological replicates). Two-tailed unpaired t-test was used for statistical analysis. (ns, no significant; *, p < 0.05; **, p < 0.01; ***, p < 0.005; ****, p < 0.001). Figure 3—source data 1. Detailed information on Co-Up and Co-Down acetylated proteins and KEGG analysis for Figure 3A. Figure 3—source data 2. PykF enzyme activity for Figure 3C, E, G, I. Figure 3—source data 3. Uncropped blots for Figure 3F, H.

To further validate the effects of protein acetylation on bacterial mobility and energy metabolism, we further detected bacterial mobility and ATP levels. The agarose exercise analysis showed that the bacterial motilities of the three antibiotic-resistant strains were weaker than that of WT (Figure 3B). Meanwhile, The ATP content of the three antibiotic-resistant strains was significantly lower than that of WT (Figure 3C). The increase of acetylation level of the corresponding proteins will lead to the increase of motility and decrease of energy level of antibiotic-resistant strains. We suggest that acetylation negatively reduces the metabolic activity of *E. coli* to maintain antibiotic resistance.

Subsequently, we found that two key acetylated rate-limiting enzymes in glycolysis, Glk and PykF, were evidently up-regulated in the three resistant strains, which were almost not detected in WT (Figure 3A,D). Among them, PykF, the key upstream kinase catalyses the reaction to produce ATP and pyruvate, the latter is the substrate for the formation of Ac-CoA, which affects the TCA cycle (Bledig et al., 1996; Zhu et al., 2008). Therefore, PykF plays an important role in the process of energy metabolism. We further tested the *in vitro* PykF enzyme activity of the WT and three resistant strains and found that PykF activity of the WT was significantly higher than that of the other antibiotic-resistant strains (Figure 3E), indicating that PykF with high acetylation levels has a lower enzyme activity.

### PykF is reversibly acetylated by CobB and PatZ

To unravel the role of acetylation of PykF in bacterial resistance to antibiotics, we further investigated the acetylation and deacetylation processes of PykF in *E.coli* which has not been reported before. A previous study reported that protein acetylation in *E. coli* is mainly catalysed by non-enzyme acetyl phosphate (AcP) and peptidyl-lysine N-acetyltransferase (PatZ) (Weinert et al., 2013). In order to study the acetylation mechanism of PykF, we expressed and purified the 6×His-tagged PykF protein and incubated it with AcP. The immunoblot with α-AcK antibody and the detection of PK enzyme activity indicated that the purified PykF protein can be acetylated with the catalysis of non-enzyme AcP. Moreover, with the extension of the incubation time, the acetylation intensity of PykF increased and its enzyme activity decreased correspondingly (Figure 3—figure supplement 1). This result indicates that non-enzyme AcP can acetylate PykF in *E. coli*, resulting in a decrease in PykF enzyme activity.

Next, CobB and PatZ were also expressed and purified and then incubated with PykF to detect their acetylation levels. The immunoblot results showed that PykF could be acetylated by PatZ in the presence of Ac-CoA (Figure 3F). Moreover, the increased acetylation level of PykF was accompanied with decreased enzyme activity (Figure 3G). In addition, when PykF was incubated with CobB and cofactor NAD+, PykF could be deacetylated by CobB. Meanwhile, when the acetylation level of PykF decreased, its enzyme activity increased accordingly(Figure 3K). However, in the presence of NAM (deacetylase inhibitors), PykF acetylation was not affected by CobB (Figure 3H, I), which further indicated that PykF is a substrate of CobB. The understanding of acetylation and deacetylation processes of PykF is conducive to the investigation of contribution of PykF acetylation in regulation of energy metabolism and bacterial resistance against antibiotics.

### K413 is the key acetylation site of PykF

To determine which lysine residues are acetylated in PykF that affect the main acetylation level changes, the MS data was analysed and three acetylation sites were found to be significantly up-regulated in the three resistant strains, including K76, K413, and K445. Among them, K413 was highly conserved in almost all the bacterial species (Figure 4A). To further test the role of K76, K413, and K445 in PykF acetylation mediated by PatZ, these lysine residues were mutated to arginine (R) or glutamine (Q). The substitution of lysine to arginine avoids acetylation but retains the positive charge, thus mimicking the non-acetylated form, while the substitution to glutamine mimics constitutive acetylation by neutralising the positive charge (Luo et al., 2004).Western blotting showed that when K413 was mutated to glutamine, the acetylation level of PykF did not increase (Figure 4B), but when K76 and K445 were mutated to glutamine, PykF acetylation was enhanced in the presence of PatZ (Figure 4C, D). These results indicate that K413 is a key acetylation site in PykF. So, we speculated that K413 contributes to PykF activity via affecting the acetylation level in resitant bacteria.

**Figure 4.**
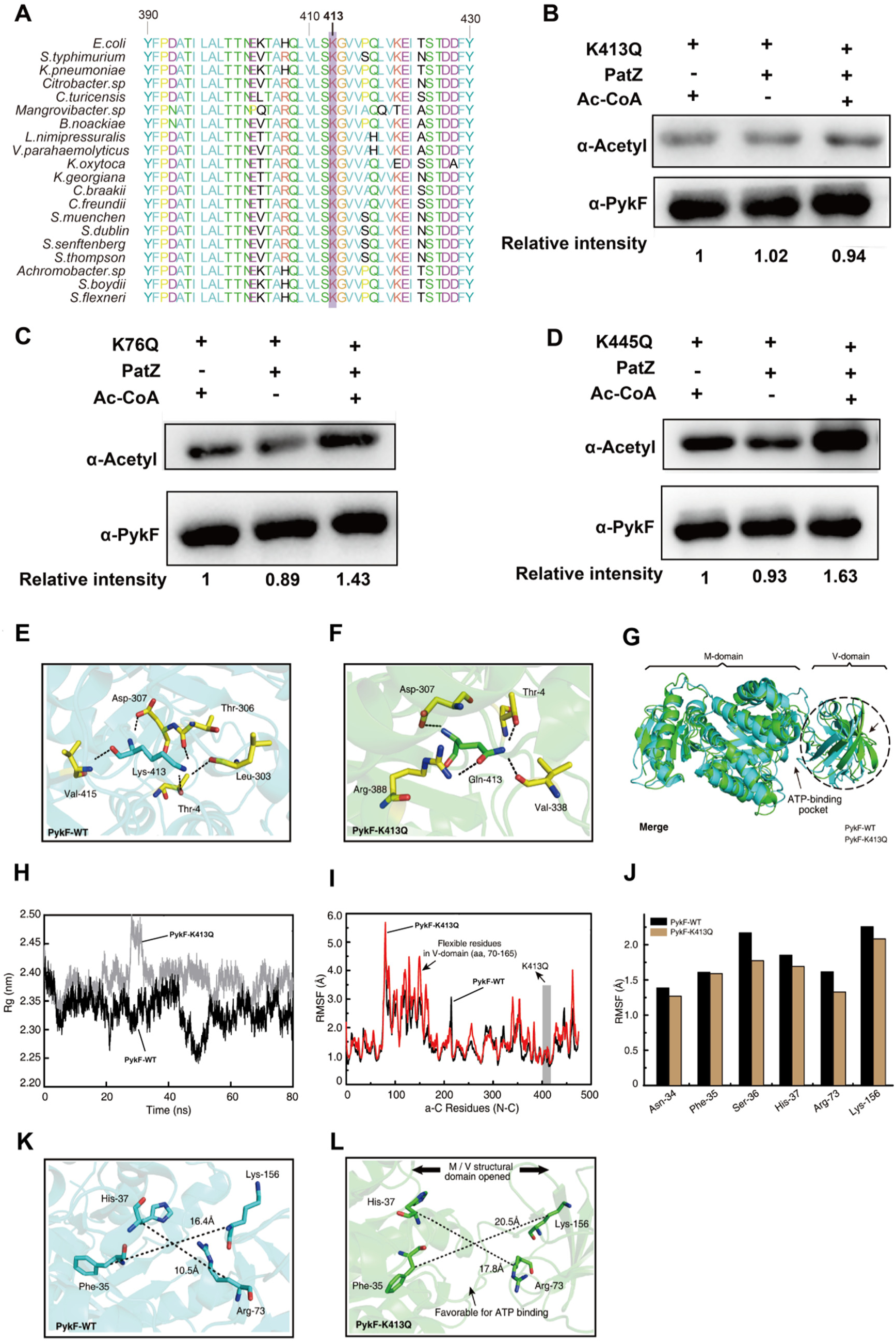
K413 of PykF is the acetylation site by PatZ. (A) Conservation analysis of PykF K413 of *Escherichia coli* through sequence alignment. Purple frame denotes the conserved lysine residues, and the result was analysed by BioEdit 7.0. (B-D) PatZ acetylates K413Q, K76Q and K445Q *in vitro*. Experiments were performed in triplicate. (E) The hydrogen bonds between Lys-413 and the surrounding residues within 3.5 Å. (F) Hydrogen bonds between Gln-413 and its surrounding residues. (G) The overall display of protein structure includes two domains (M-domain and V-domain). Comparison of the average conformation of PykF-WT and PykF-K413Q in equilibrium state. The dashed box represents the three-dimensional structure area with the largest conformational difference. The active centre of ATP is marked by a black arrow. (H) Time evolution of the C-alpha of the Rg values. (I) The RMSF values of the C-alpha for all residues of PykF-WT and PykF-K413Q proteins calculated over the 80-ns trajectory. The RMSF value of residues in V-domain of PykF-K413Q are higher than that of PykF-WT (black arrow). (J) The RMSF of the C-alpha profile of residues in the ATP active binding sites of PykF-WT and PykF-K413Q MD trajectories. (K-L) The differences of the ATP-binding pocket located between the V- and M-domains in PykF-WT or PykF-K413Q were compared, and the distances between the selected residues were measured. Figure 4—source data 1. Uncropped blots for Figure 4B, C, D.

To further investigate the effects of Lys-413 acetylation on PykF conformation and enzyme activity, we analysed the conformational differences of PykF-WT and PykF-K413Q in equilibrium state using molecular dynamics (MD) simulation. The results showed five hydrogen bonds formed between Lys-413 and the surrounding residues Thr-4/Leu-303/Thr-306/Asp-307/Val-415 (Figure 4E) in PykF, while Gln-413 interacted with Thr-4/Asp-307/Val-338/Arg-388 in PykF-K413Q (Figure 4F), which means that the replacement of Lys-413 by Gln induced the local microenvironment change of the key acetylated site. By observing the whole conformation of PykF-WT, we found that about 96 residues (aa, 70-165) formed a single Vice-domain (V-domain), while the remaining residues formed the main-domain (M-domain) (Figure 4G). These two domains jointly contribute to the biological activity of pyruvate kinase. According to the description of this protein in the UniProt database, the ATP pocket is located at the interface between the M- and V-domains. We merged the conformations of PykF-WT and PykF-K413Q under equilibrium state, and found that the overall conformation of the V-domain deviated from that of the M-domain in the mutant protein, which was conducive to the ATP pocket showing opened state, facilitating ATP binding. However, ATP inhibits its activity as PykF in *E. coli* is a type I isozyme (Fenton & Hutchinson, 2009; Mattevi et al., 1995; Valentini et al., 1995). Based on the results of MD simulation, we found that the Rg values of C-alpha of PykF-K413Q were larger than that of PykF-WT (Figure 4H), which indicates that the replacement of Lys-413 by simulating Gln may induce conformational looseness of the whole protein. Moreover, the residues of V-domain of PykF-K413Q showed higher RMSF values than that of PykF-WT (Figure 4I), indicating the residues of V-domain are more flexible in the mutant protein. We further calculated the RMSF values of the typical ATP-binding site residues Asn-34/Phe-35/Ser-36/His-37/Arg-73/Lys-156 of PykF and found that they were more stable in PykF-K413Q than that in PykF-WT (Figure 4J). In addition, the induced conformation dramatically increased the distance among the ATP-binding site residues in PykF, specifically Phe-35 and Lys-156, and His-37 and Arg-73 (Figure 4K, L), resulting in larger solvent accessible surface area of ATP active centre and longer distance between M- and V-domains which jointly contribute to ATP binding. These results suggested that all these conformational changes induced by acetylation may promote the binding of PykF-K413Q with ATP, which will further inhibit the enzyme activity.

### Mimic deacetylation of PykF K413 reduces bacterial resistance ability

As we know, PykF is an important key rate-limiting enzyme for the ATP production during glycolysis which provides pyruvate for the TCA cycle(Bledig et al., 1996). Interestingly, we identified K413 acetylation of PykF in all the three resistant strains, but not in WT. So, we speculated that K413 acetylation could contribute to the maintenance of resistance in bacteria. In order to further study whether the deacetylation of K413 site affects drug resistance, we knocked out the *pyk*F gene in the three resistant strains to construct Δ*pykF*-Amp/kan/pol mutant strains, and then introduced the plasmid with wild type PykF, PykF-K413Q, and PykF-K413R gene into cells to get eWT-Amp/Kan/Pol, eK413Q-Amp/Kan/Pol and eK413R-Amp/Kan/Pol strains, respectively. At the same time, in order to make the experiment more clinically meaningful, we used the same method to knock out the *pykF* gene in a clinical multi-drug-resistant *Escherichia coli*, (clinical strain number 966113) (CMR) and then introduced the above mentioned three plasmids into it and named as eWT-CMR, eK413Q-CMR, and eK413R-CMR. We collected the constructed bacteria at OD_600_ of 0.8, and detected their PykF enzyme activity and ATP content in cells. The results showed that in the four resistant strains, K413 mimic deacetylation resulted in an increase in PykF enzyme activity (Figure 5A-D) and ATP content (Figure 5E-H). WB detection showed that the expression level of PykF in three complementary strains of each kind of resistant strains were similar (Figure 5—figure supplement 1A-B). Importantly, it was found that the supplement with PykF-K413R resulted in the decrease of the antibiotic resistance of the three resistant strains domesticated in the laboratory (Fig 5I-K) and CMR strains to ampicillin (Figure 5L). In addition, we also carried out the corresponding experiments in the sensitive *E. coli* BW25113 (Figure 5M, N) and obtained the same conclusion as above. Otherwise, we directly expressed WT PykF, PykF K413R and PykF K413Q in sensitive *Salmonella* ATCC14028 and detected their antibiotic resistance, and also found that f PykF-K413R contribued *Salmonella* resistance (Figure 5O, P). These results indicate that K413 is an important lysine acetylation site, whose deacetylation will increase PykF activity and ATP production, correlating with the decrease of bacterial resistance to antibiotics.

**Figure 5.**
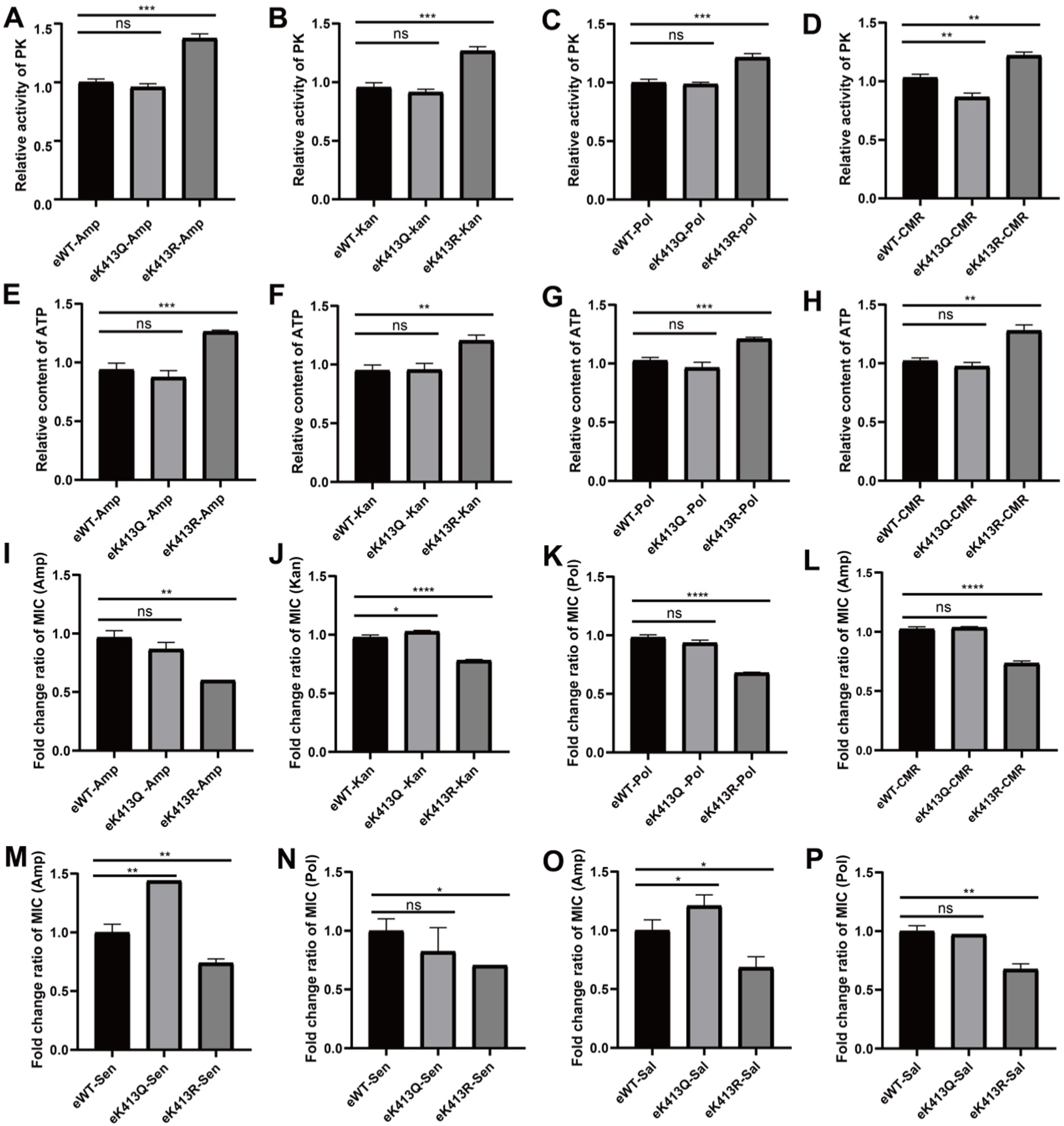
Deacetylation of PykF K413 reduces bacterial resistance. (A-D) PykF enzyme activity detection, (E-H) ATP content detection, (I-L) MIC determination of the resistance strains in which *pykF* was knocked out and then complemented with wild-type *pykF*, K413Q and K413R plasmid.(M-N) MIC change of sensitive *E.coli* BW25113 with *pykF* gene knocked out. (O-P) MIC changes of ampicillin and polymyxin after supplementation of wild-type PykF and mutant PykF in *Salmonella*. (A-P) (n = 3 independent biological replicates).Two-tailed unpaired t-test (Welch’s t test) was used for statistical analysis. (ns, no significant; *, p < 0.05; **, p < 0.01; ***, p < 0.005; ****, p < 0.001). Figure 5—source data 1. PykF enzyme activity, ATP content and MIC for Figure 5.

In summary, with immune enrichment and DIA-based quantitative proteomics, we found a common regulatory mechanism of antibiotic resistance by acetylation in bacteria. Lysine acetylation negatively regulates energy metabolism and positively regulates bacterial motility, which reduces the overall metabolic level of bacteria to maintain drug resistance. The key limiting-rate kinase PykF in glycolysis was highly acetylated in the three resistant strains and K413 was found to be an important acetylation site for the enzyme activity. Change in the acetylation state to reduce the level of PykF K413 acetylation increases bacterial energy metabolism and the sensitivity of bacteria to antibiotics. So, this study uncovers a new systematic regulatory mechanism of drug-resistant strains (Figure 6) and provides novel insights for the development of new antibacterial drugs.

**Figure 6.**
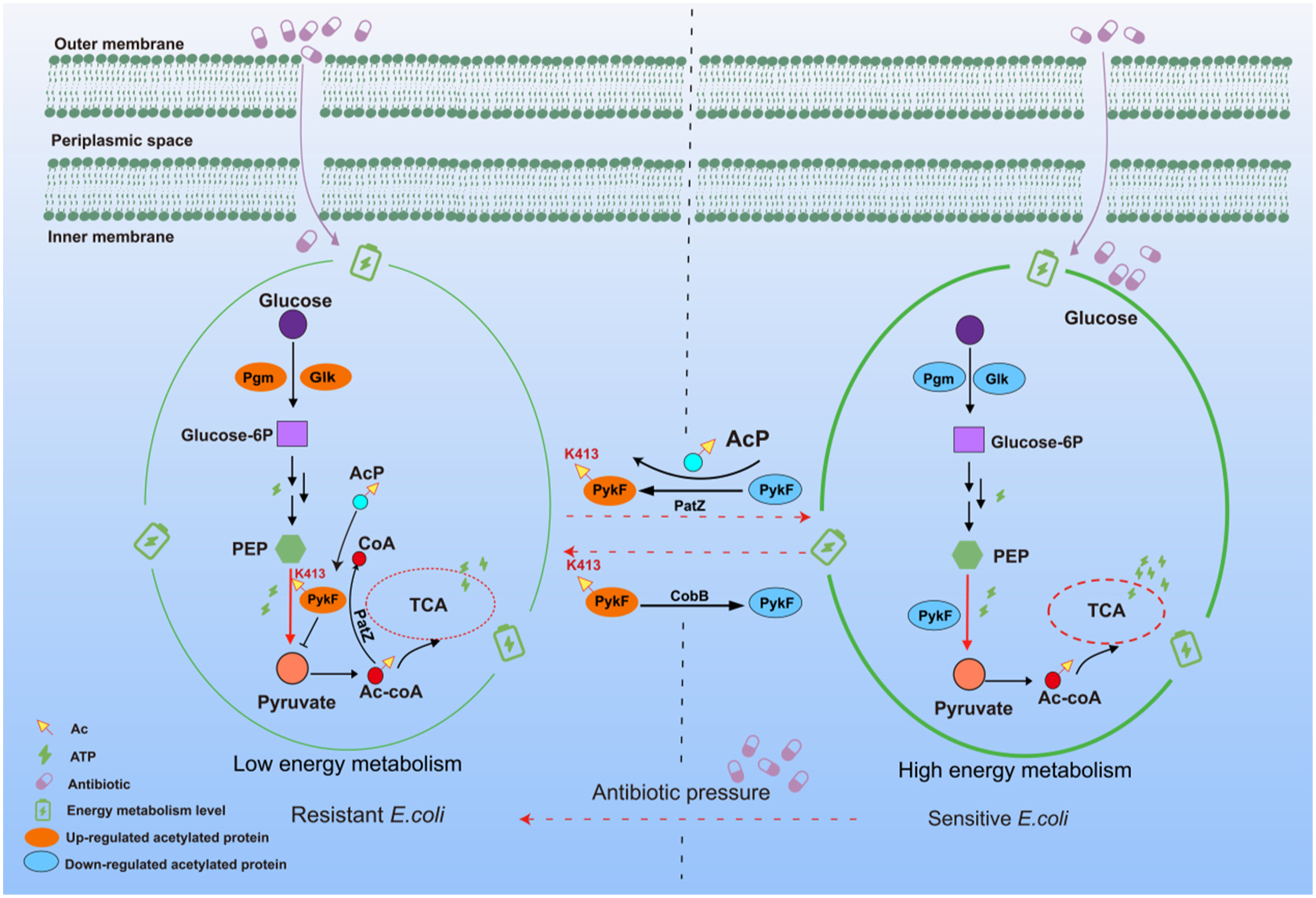
Illustration of lysine acetylation regulating bacterial resistance. Under the long-term pressure of antibiotics, *E. coli* acquired drug resistance through acetylation negative regulation of its own metabolism. In drug-resistant bacteria, it was found that the acetylation of lysine at position 413th of pyruvate kinase PykF in the pyruvate cycle decreased its enzyme activity and further inhibited the process of TCA cycle to maintain low energy metabolism. In this study, it was found that deacetylation of PykF K413 could restore the sensitivity of *E. coli* to antibiotics (Kanamycin, ampicillin, polymyxin B) and make it obtain higher energy metabolism.

## DISCUSSION

Over the past decades, the abuse of antimicrobial drugs has led to the appearance of various super-resistant and multi-resistant strains which have seriously endangered human health(Hidron et al., 2009; Ling et al., 2006). Therefore, in the recent years, the prevention and treatment of infection caused by drug-resistant strains and the rational use of antibiotics have received worldwide attention . A systematic study of the mechanisms of bacterial resistance is necessary to combat the infections caused by drug-resistant strains. The development of bacterial resistance is a process in which multiple mechanisms work together(Baron et al., 2014; Hidron et al., 2009; Ling et al., 2006; Thongboonkerd, 2010). Under the pressure of antibiotics, bacteria can gradually adapt to the environment by regulating their own physiological state, thus obtaining drug resistance(Lin et al., 2014; Lobritz et al., 2015). Meanwhile, more and more studies have pointed out that change in bacterial metabolic level can significantly affect the bactericidal effect of antibiotics(Su et al., 2018). Post-translational modifications of proteins are involved in many important metabolic pathways including the regulation of key catalytic enzymes.

We conducted quantitative acetylation proteomics research to explore the relationship between acetylation and bacterial resistance. It is noteworthy that certain specific acetylated proteins were only detected in one kind of resistant strains, suggesting that different antibiotic-resistant strains may have specific acetylated proteins involved in the regulation of resistance. After statistical analysis of these unique acetylated proteins, 13 specific acetylated proteins were found to be involved in the resistance regulation of ampicillin-resistant strains, 28 were observed in kanamycin-resistant strains, and 58 were involved in polymyxin B-resistant strains. According to different biological processes, specific acetylated proteins that may be related to different antibiotic resistances were determined (Table 1). For example, acetylated entericidin B (EcnB), a membrane-associated protein, was only identified in ampicillin-resistant strains. EcnB plays a role in bacteriolysis and affects the synthesis of membrane proteins. Changes in membrane protein structure might affect the efficacy of ampicillin [6], which indicates that EcnB acetylation may be uniquely involved in the regulation of ampicillin resistance. In addition, the unique acetylated protein AtoC, member of the two-component regulatory system AtoS/AtoC(Lee et al., 2011), was only found in kanamycin-resistant strains. AtoC not only acts as a transcriptional regulator, but also acts as a post-translational regulator(Theodorou et al., 2012; Theodorou et al., 2007). It can inhibit the biosynthesis of polyamines by regulating ornithine decarboxylase. However, as the target site of kanamycin is the 30S subunit, we speculated that AtoC acetylation might inhibit the biosynthesis of polyamines, inhibiting the complete structure formation of the 30S subunit. This change in the target site might lead to the bacterial resistance against kanamycin. Besides, in polymyxin B-resistant strains, we found that the unique acetylated proteins ArnA and ArnB, modify lipid A to change the target site of polymyxin B attack(Breazeale et al., 2005; Breazeale et al., 2002). Therefore, these two unique acetylated proteins may be involved in regulating the bacterial resistance to polymyxin B. These highly acetylated proteins indicate that different resistant strains may have certain distinct regulatory mechanisms. Moreover, the above results indicate that these proteins may be used as biomarkers to identify different types of resistant strains, but further research and verification are still needed.

**Table 1.** Specific acetyl proteins identified in different resistance strains.

| Specific acetylated protein related to ampicillin resistance |  |
| --- | --- |
| Characteristics | Acetylated proteins and sites |
| Multidrug resistance protein | MdIA K232 |
| Membrane lipoprotein | EcnB K45 |
| DNA binding protein | DecR K115, YrdD K123 |
| Stimulus sensor | InaA K176 |
| Glucose dehydrogenase | Gcd K274/K685 |
| Sugar phosphotransferase system | YadI K82 |
| Specific acetylated protein related to kanamycin resistance |  |
| Characteristics | Acetylated proteins and sites |
| Two-component system | AtoC K126/K127, NarL K100 |
| ABC transporter | HisJ K120 |
| Kinase | NanK K197 |
| Membrant protein | YejM K220, TolA K10, MinC K26 |
| Toxin-antitoxin system | DinJ K73 |
| Specific acetylated protein related to polymyxin resistance |  |

| Characteristics | Acetylated proteins and sites |
| --- | --- |
| Polymyxin resistance protein | ArnA K181/K422/K442/K458/K532/K611, ArnB |
| LPS related enzymes | Ais K76/K188, EptC K9/K547 |
| Toxin-antitoxin system | YhaV K73 |
| Kinase | Tmk K4/K209, PdxY K108/K120 |
| Two-component system | BasR K164, UvrY K41, YehU K327 |
| ABC transporter | HisP K148 |
| DNA binding protein | Ssb K88, MutY K103, SlmA K179, StpA K107 |
| Reductase | Fre K83, ProA K15, ProC K226, YbjS K334 |
| Iron metabolism related protein | EntC K237, Fes K47, IscU K46, IscA K89, MenF K120 |

More importantly, bioinformatics analysis showed a common characteristic in three resistant strains that acetylation can regulate *E. coli* metabolism in two ways to maintain antibiotic resistance. Firstly, acetylation positively regulates bacterial motility by reducing the acetylation level of the flagellar movement-related proteins, among which FliA and FlhD were identified among the Co-Down-regulated DAPs. FlhD is an upstream transcription factor that can form a transcription activation complex FlhD4C2 with FlhC, and FlhD integrates a large number of input signals from global regulatory factors and determines cellular mobility (Chilcott & Hughes, 2000; Lee et al., 2011). In addition, FlhD can regulate the expression of FliA which significantly affects the biofilm formation ability (Duan et al., 2013), indicating that mobility is also an important factor affecting the biofilm formation. Moreover, the biofilm will further contribute to bacterial resistance. Therefore, we believed that the decrease in acetylation levels of FlhD and FliA proteins may promote the development of bacterial resistance. However, the mechanism behind specific control needs further verification. Secondly, acetylation negatively regulates the energy metabolism of bacteria by increasing the level of acetylation of central metabolism-related proteins. Previous studies have shown that regulation of the bacterial metabolic level can improve the resistance efficacy of antibiotics. For example, the addition of exogenous glutamic acid can increase the sensitivity of bacteria to aminoglycoside antibiotics (Su et al., 2018), but the related metabolic regulation mechanism is still unclear. Therefore, from the point of view of lysine acetylation, we compared the WT with the resistant strains to explore the similarities in resistance mechanisms. The Co-Up of DAPs was significantly enriched in the central metabolic regulation process, which is consistent with the previously reported acetylation concentrated in the carbon metabolism regulation process(Castano-Cerezo et al., 2014; Weinert et al., 2013; J. M. Zhang et al., 2009).

In addition, the pyruvate cycle (oxaloacetate-PEP-pyruvate-acetyl-CoA) can produce a lot of energy. When the enzymes in the pyruvate cycle are inhibited or genes are inactivated, the TCA cycle is blocked (Su et al., 2018). PykF is the key kinase in the pyruvate cycle, which is essential for the production of ATP and pyruvate, and affects the level of energy metabolism in bacteria (Su et al., 2018). Our study found that acetylated form of PykF was significantly up-regulated in the three drug-resistant strains. Moreover, our further studies found that its enzyme activity is negatively regulated by acetylation, and the K413 site is highly conserved and key acetylation site. Deacetylation at this site can increase the enzymatic activity of PykF, which leads to an increase in the energy metabolism of bacteria, which in turn increases the sensitivity of drug-resistant strains to antibiotics. The same was observed even in the clinical multiple resistant strains to a certain level. However, compared with the sensitive bacteria, the sensitivity to antibiotics cannot be fully restored, that is, it still retains a certain degree of resistance may be due to other energy compensation mechanisms of bacteria.

In conclusion, we found that lysine acetylation exhibits similarities in antibiotic-resistant strains with different resistances. By positively regulating bacterial motility and negatively regulating energy metabolism, the metabolic level of bacteria is reduced to maintain antibiotic resistance. Furthermore, our research found that deacetylation of the K413 site of the PykF protein can increase the enzyme activity of PykF, leading to an increase in the energy metabolism of bacteria, which in turn increases the sensitivity of antibiotic-resistant strains to antibiotics. The metabolic network of lysine acetylation to regulate bacterial resistance have been described here and key regulatory proteins were detected, which provide a new perspective for studying bacterial resistance mechanisms.

## MATERIALS AND METHODS

### Detection of bacterial rresistance to antibiotics

All the bacterial strains, plasmids, and primers used in this study are listed in the Table 2-3. The minimum inhibitory concentration (MIC) for Amp was determined via microdilution in a 48-well plate. The bacteria were diluted to an OD_600_ of 0.05 in Luria-Bertani (LB) broth, and then 500 μL of diluted culture was added to each well of a 48-well plate containing Amp at different concentrations, including 2, 4, 6, 8, 10, 12 and 14 μg/mL. Cultures were incubated at 37 °C for 24 h and then growth was detected using a microplate reader. The drug concentration in which the bacterial growth of an OD_600_ below 0.1 was observed was recorded as the MIC(Andrews, 2001). The MIC of WT *E. coli* BW25113 was experimentally determined as 10 μg/mL. Three single colonies of WT *E. coli* BW25113 were cultured overnight and then diluted 1:100 in fresh LB medium with or without (5 μg/mL) of Amp. After 24 h of incubation at 37 °C with agitation, the MIC was determined again. Then, the bacteria are continuously subcultured at subMIC, and the concentration of antibiotics is gradually increased to improve the resistance of bacteria. The same procedure was repeated until the MIC of Amp increased to 481.7 μg/mL, being 60-fold of that of the original MIC. Domestic strain of *E. coli* was also tested with kanamycin and polymyxin B. The resistance development of Δ*patZ* and Δ*cobB* strains against ampicillin and polymyxin B were also tested with the method mentioned above.

**Table 2.** Strains and plasmids used in this study.

| Strains and plasmids | characteristics | Sources or references |
| --- | --- | --- |
| **Strains** |  |  |
| *E. coli* BL21 | Expression strain | Laboratory stock |
| *E. coli* BW25113 | Wild type *Escherichia coli* strain | Laboratory stock |
| *Salmonella* ATCC14028 | Wild type *Salmonella* strain | Laboratory stock |
| Re-Amp | Ampicillin resistant *Escherichia coli* strain | This study |
| Re-Kan | Kanamycin resistant *Escherichia coli* strain | This study |
| Re-Pol | Polymyxin B resistant *Escherichia coli* strain | This study |
| $\Delta pykF$ | Wild type *Escherichia coli* BW25113 strain $\Delta pykF$ | NBRP (NIG, Japan)<sup>35</sup> |
| $\Delta cobB$ | Wild type Escherichia coli BW25113 strain $\Delta cobB$ | NBRP (NIG, Japan)<sup>35</sup> |
| $\Delta patZ$ | Wild type Escherichia coli BW25113 strain $\Delta patZ$ | NBRP (NIG, Japan)<sup>35</sup> |
| $\Delta pykF:: Re-Amp$ | Ampicillin resistant Escherichia coli strain $\Delta pykF$ | This study |
| $\Delta pykF:: Re-Kan$ | Kanamycin resistant Escherichia coli strain $\Delta pykF$ | This study |
| $\Delta pykF:: Re-Pol$ | Polymyxin B resistant Escherichia coli strain $\Delta pykF$ | This study |
| $\Delta pykF:: CMR$ | Multidrug resistant Escherichia coli strain $\Delta pykF$ | This study |
| eWT-Amp | PBSU101 with *PykF*, overexpressed in $\Delta pykF:: Re-Amp$ | This study |
| eWT-Kan | PBSU101 with *PykF*, overexpressed in $\Delta pykF:: Re-Kan$ | This study |
| eWT-Pol | PBSU101 with *PykF*, overexpressed in $\Delta pykF:: Re-Pol$ | This study |
| eWT-Sen | PBSU101 with *PykF*, overexpressed in $\Delta pykF$ | This study |
| eWT-Sal | PBSU101 with *PykF*, overexpressed in *Salmonella* | This study |
| eWT-CMR | PBSU101 with *PykF*, overexpressed in $\Delta pykF:: CMR$ | This study |
| eK413Q -Amp | PBSU101 with *PykF* (K413Q), overexpressed in $\Delta pykF:: Re-Amp$ | This study |
| eK413Q-Kan | PBSU101 with *PykF* (K413Q), overexpressed in $\Delta pykF:: Re-Kan$ | This study |
| eK413Q-Pol | PBSU101 with *PykF* (K413Q), overexpressed in $\Delta pykF:: Re-Pol$ | This study |
| eK413Q-Sen | PBSU101 with *PykF* (K413Q), overexpressed in $\Delta pykF$ | This study |
| eK413Q-Sal | PBSU101 with *PykF* (K413Q), overexpressed in *Salmonella* | This study |
| eK413Q-CMR | PBSU101 with *PykF* (K413Q), overexpressed in $\Delta pykF:: CMR$ | This study |
| eK413R-Amp | PBSU101 with *PykF* (K413R), overexpressed in $\Delta pykF:: Re-Amp$ | This study |
| eK413R-Kan | PBSU101 with *PykF* (K413R), overexpressed in $\Delta pykF:: Re-Kan$ | This study |
| eK413R-Pol | PBSU101 with *PykF* (K413R), overexpressed in $\Delta pykF:: Re-Pol$ | This study |
| eK413R-Sen | PBSU101 with *PykF* (K413R), overexpressed in $\Delta pykF$ | This study |
| eK413R-Sal | PBSU101 with *PykF* (K413R), overexpressed in *Salmonella* | This study |
| eK413R-CMR | PBSU101 with *PykF* (K413R), overexpressed in $\Delta pykF:: CMR$ | This study |
| **Plasmids** |  |  |
| pET28b | Expression vector | Laboratory stock |
| pET28b-*pykF* | *pykF* in pET28b | This study |
| pET28b-*pykF* (K413Q) | *pykF* (K413Q) in pET28b | This study |
| pET28b-*pykF* (K413R) | *pykF* (K413R) in pET28b | This study |
| PBSU101-*pykF* | *pykF* in PBSU101 | This study |
| PBSU101-*pykF* (K413Q) | *pykF* (K413Q) in PBSU101 | This study |
| PBSU101-*pykF* (K413R) | *pykF* (K413R) in PBSU101 | This study |
| pET28b-*cobB* | *cobB* in pET28b | This study |
| pET28b-*patZ* | *patZ* in pET28b | This study |

**Table 3.**
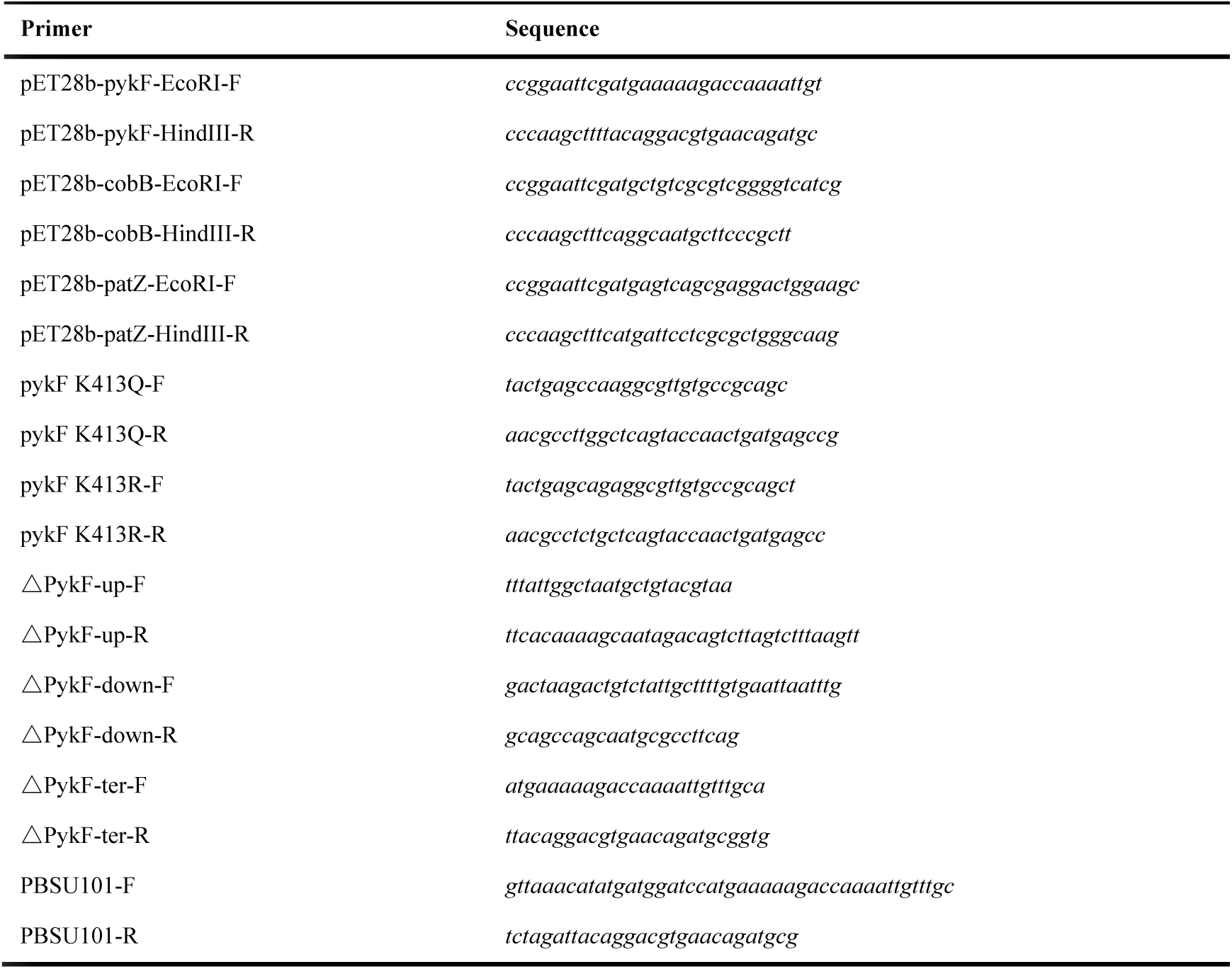
Primers and peptides used in this study.

### Cell culture and protein exaction

All strains used herein were activated in LB medium at 37 °C overnight, transferred into fresh LB medium (1:100, v/v) for culturing, and harvested after the OD_600_ approached 0.8 through centrifugation at 8000 g for 10 min at 4 °C. The cell pellets were washed three times with ice-cold PBS and then sonicated in 500 μL of SDS lysis buffer supplemented with protease inhibitor cocktail (Roche, Basel, Switzerland), 1 mM phenylmethanesulfonyl fluoride (PMSF), and deacetylase inhibitor mix (Beyotime, Nanjing, China). After centrifugation at 12000 g for 30 min, the supernatant was collected. The protein concentration was determined with the BCA Protein Assay Kit (Thermo Fisher Scientific, Shanghai, China).

### Western blotting

The protein is loaded into the 12% SDS-PAGE gel from each sample and then transferred to the polyvinylidene fluoride (PVDF) membrane (Milliport, USA). Acetylated antibody (PTM BioLab, PTM-101) was incubated with polyvinylidene fluoride membrane at 4 °C, and horseradish peroxidase-conjugated goat anti-mouse was used as secondary antibody. The results were visualized with Clarity Western ECL Substrate (Bio-Rad, USA) and captured with ImageMaster2D Platinum 6.0 (GE Healthcare, USA). Meanwhile, the SDS-PAGE gel stained with Coomassie brilliant blue R250 was used as the load control.

### Protein digestion and affinity enrichment of acetylated peptides

The protein digestion and affinity enrichment of the acetylated peptides were performed as described in a study conducted by us previously (Liu et al., 2018). Briefly, the protein extract was reduced with 8 M urea and 50 mM dithiothreitol (DTT, 37°C, 1 h) and alkylated with 100 mM iodoacetamide (IAA, 25 °C, 30 min). Samples were transferred into the 30 kDa ultracentrifugal filters (Sartorius Stedim Biotech, Shanghai, China) and washed thrice with 8 M urea and 50 mM triethylammonium bicarbonate buffer (TEAB). Protein and trypsin were mixed at a mass ratio of 30:1 for digestion at 37 °C for 16 h. Then, the peptides were lyophilised at -80°C for further analysis. The α-AcK antibody beaded agarose kit (PTM biolabs, Hangzhou, China) was used for enriching acetylated peptides. The lyophilised peptide was dissolved in 200 μL of pre-cold IP Buffer, then gently mixed with antibody conjugated beads. The ratio of peptides to beads was 3 mg peptides to 30μL of drained antibody beads. After incubating at 4°C overnight, the peptides were washed with wash buffer I and wash buffer II thrice and the acetylated peptides were eluted with 100 μL of elution buffer. Finally, the elutes were dried for MS analysis.

### Data-independent acquisition by MS

Peptides were dissolved in 0.1% formic acid and diluted to 0.5 μg/μL. To build the precursor ion library, 1 μL of each sample was taken to prepare a mixed sample. IRT-standard provided by the iRT-Kit (Biognosys, Schlieren, Switzerland) at 1/10 by volume was added to all the samples. The mixed sample was analysed thrice by Orbitrap Fusion Lumos (Thermo Fisher Scientific) in data dependent acquisition (DDA) mode. The DDA library building parameters were set as follows: ion source type: NSI, positive ion reflection mode; ion transfer tube temperature: 320°C; pressure mode: standard; default charge state: 2; do data dependent experiment if no target species are found: False; MSn Level: 1; detection type: ion trap; resolution: 60K; mass range: normal; scanning range: 400-1500 m/z; maximum injection time: 50 ms; mass tolerance low: 10; mass tolerance high: 10 ; filter type: intensity threshold; signal strength: 50000; separation mode: quadrupole; activation type: HCD; and collision energy: 30%.

Next, each individual sample was analysed in the data-independent acquisition (DIA) mode with the same instrument. Three biological replicates were performed for each sample. The DIA parameter setting was basically the same as that of the DDA analysis except for the following: MS scan range, 350–1200 m/z; MS/MS scan range, 200–2000 m/z.

### Data processing

To build a DDA library, the following command was used to search the original DDA file: Customise the Sequest HT (v2.5) engine in Proteome Discoverer software version 2.1 (Thermo Fisher Scientific) with the Uniprot-*E. coli* K12 FASTA database and iRT standard peptide sequence customisation. Compare with reference database in FASTA format, the search parameters were set as follows: MS tolerance: 10 ppm; fragment mass tolerance is 0.02 Da; enzyme: trypsin; static modification: carbamidomethylation (C); dynamic modification: oxidation of methionine, deamination of Q and N, N-terminal Acetyl and acetylation of K. Filter peptides to obtain high confidence (FDR, 1%) with a minimum peptide length of seven aa. The selected protein meets the following conditions: (1) protein level FDR ≤ 1%; (2) unique peptides ≥ 2. The pdResult file searched by DDA was exported from PD software and imported into Spectronaut software version 10 (Biognosys) to construct an ion spectrometry library.

Next, we used the following command to convert the DIA original file to htrm format: HTRMS converter provided by Spectronaut. Finally, the DIA htrm file, the DDA original file, the DDA pdResult file, and the customised Uniprot-*E. coli* K12 database + iRT standard peptide FASTA file were loaded into Spectronaut, and processed through the BGS factory settings. Basically, the default parameters were used, except the enzyme Modify from trypsin/P to trypsin and add acetylation of K. Protein was inferred by software, standard q value of 0.01 was used to obtain the quantitative information at protein level, which was used for subsequent analysis.

### Bioinformatics analysis

In order to obtain the actual change intensity of acetylation, we used the following formula to deduct the protein background: (A/a)/(B/b), where ‘A’ represents the quantitative acetylation value of resistant strains, ‘a’ represents the quantitative protein of resistant strains, ‘B’ represents the WT strain quantitative acetylation value, and ‘b’ represents the quantitative protein value of the WT strain. In this way, we obtained the relative up-down fold change relative to that of the WT strain and the subsequent data analysis.

Gene Ontology (GO) enrichment analysis was performed in Blast2GO(Conesa et al., 2005) with Fisher’s exact test of FDR < 0.05. Kyoto Encyclopedia of Genes and Genomes (KEGG) pathway enrichment analysis was performed in the “Wu Kong” platform. Functional protein domains were predicted with Pfam 31.0 database(Finn et al., 2016). “Wu Kong” platform was also used for the analysis of amino acid sequences and generation of the sequence logos.

### Cloning, expression, and purification of target proteins

According to the method previously described(Miao et al., 2018; Zheng et al., 2020), *PykF*, *cobB and patZ* genes were amplified from the genomic DNA of *Escherichia coli* BW25113 (primers are listed in Table 3). The PCR product and pET28b plasmid were digested with restriction endonuclease EcoRI and HindIII (Japanese Takara), then ligated with T4 DNA Ligase (Japanese Takara), and transformed into Escherichia coli BL21. The strains verified by sequencing were cultured in LB medium containing 50 μg/mL kanamycin at 37 °C. When OD600 reached 0.6, 0.5 mM isopropyl-D-thiogalactoside (IPTG; Sigma) was added and cultured for 5 h to induce protein expression. The bacteria were centrifuged for 8000g at 4 °C and washed with 0.01M phosphate buffered saline (PBS) for 3 times. The collected cells were crushed with 0.01M PBS suspension and cracked under high pressure. And the supernatant was obtained by centrifugation and 30min at 12000g. Then, the recombinant protein was purified by Ni-NTA affinity chromatography column (QIAGEN).

### Generation of gene-deletion mutants

Knockout of pykF is accomplished by inactivating chromosomal genes(Baba et al., 2006). To construct these mutants, primers for cloning homologous pykF are listed in Table 3. The prepared electrotransformation competent transformed pCas9 plasmid, the monoclonal LB grown at 30°C was cultured in liquid culture, the cas9 protein was induced with arabinose, the electrotransformed competent cells were prepared, the homologous fragments and psgRNA plasmids were transformed, and gentamicin was coated culture resistant plate at 30°C. The deletion mutant was confirmed by PCR. The test primers are listed in Table 3.

To construct the complement strains, the constructed pET28b-pykF, pET28b-pykF (K413Q) and pET28b-pykF (K413R) plasmid was amplified by PBSU101 primers (Table 3), and the PBSU101 plasmid was digested by BamHI and XbaI (Japanese Takara). Then PBSU101-pykF, PBSU101-pykF (K413Q) and PBSU101-pykF (K413R) were was obtained by homologous recombination (Clone ExpressII One Step Cloning Kit, Vazyme) of the amplified product and the digested linearized plasmid. Next, the obtained plasmid was introduced into the corresponding strain.

### Site-directed mutagenesis of pykF

The Mut Express II Fast Mutagenesis Kit V2 (Vazyme Biotech, C214-01) was used to introduce base substitutions into the WT *pykF* allele through the corresponding primers according to the manufacturer’s instructions. All the site-directed mutants were confirmed by DNA sequencing. All primers used for site-directed mutagenesis were listed in Table 3.

### Motility assays

The motility assays were performed as described in our previous study(Du et al., 2019). Bacterial solution (1 µL) was inoculated at the centre of the agarose plate (Swimming plate) and incubated at 37 °C for 24 h. Then, the diffusion area of different bacteria on the plate was calculated using the software Image J (National Institutes of Health, Bethesda, MD, USA).

### *In vitro* modification assay

All the *in vitro* modification assays were performed as described in the previous studies(Li et al., 2017; Ren et al., 2016). For the PatZ modification assay, PykF (0.2 μg/μl) was purified and incubated at 37°C for 6 hours in the presence or absence of PatZ(0.2 μg/μl) and Ac-CoA (0.2 mM). For the CobB deacetylation assay, PykF (0.2 μg/μl) protein was incubated at 30°C for 8 hours in the presence or absence of CobB (0.6 μg/μl), NAD^+^ (3 mM), and NAM (30 mM). For the AcP modification assay, PykF was incubated at 37°C in the presence or absence of 20 mM AcP, and the samples were collected after 0 hours and 3 hours.

### MD simulation

In this study, the MD simulations were carried out in the 80 ns trajectory by the Gromacs 2019.5 package with SPC water model in the Gromos43a1 force field. The PyMOL software was used to perform the mutation from Lys-413 to Gln-413. Next, the PDB structures of PykF-WT and PykF-K413Q were used as starting conformations in one cubic periodic box and the minimum distance between the protein and the box boundary was set at 1.0 nm. The periodic boundary condition of the MD system was suitable for the three directions of X/Y/Z. The solute (0.15 mol/L NaCl salt) was added in solvent to neutralise the system. The electrostatic interactions were computed using the particle mesh ewald (PME) method and the atomic motion was calculated using the leapfrog algorithm within the MD system. The 400 steps energy minimisation was performed using the steepest descent energy method. Next, we carried out the 25000 steps energy minimisation by the conjugate gradient method and the 50 ps position restraint simulation in each system. Each MD trajectory was carried out with a random initial velocity at a constant temperature of 300 K.

### MD trajectory analysis

The root mean square fluctuations (RMSF) of each alpha-C of the residue was calculated using gmx rmsf tool, and the radius of gyration (Rg) values of atomic evolution with the simulation time was calculated using the gmx gyrate tool of Gromacs 2019.5. The conformational changes between PykF-WT and PykF-K413Q were observed from the MD simulations trajectory by VMD-1.9.1 software. Then, the average distances of the typical ATP binding site residues of PykF-WT and PykF-K413Q, such as Phe-35 and Lys-156, His-37 and Arg-73, respectively, were measured by gmx distance. The hydrogen bonds around Lys-413 or Gln-413 were analysed using gmx hbond and PyMOL and Origin 8.5 software were used to draw the structure diagram.

### ATP and pyruvate kinase activity assays

ATP levels and pyruvate kinase activity were determined with the ATP Assay Kit (Beyotime, S0026) and Pyruvate Assay Kit (Solarbio, BC2205), respectively, according to the manufacturer’s instructions.

## ACKNOWLEDGMENTS

This work was supported by the National Natural Science Foundation of China (21977037, 21571082, to X. S..), the National Key R & D Program of China (2017YFA0505100 to Q.-Y. H.), Guangzhou Science and Technology Grant (201607010228, to X. S.).

We would like to thank Editage (www.editage.cn) for English language editing and ‘Wu Kong’ platform (https://www.omicsolution.com/wkomics/main/) for relative bioinformatics analysis. We thank Dr. Shisheng Wang (West China Hospital, Sichuan University) and Dr. Chengpin Shen (Omicsolution Co., Ltd) for giving some advices about data analysis, and Dr. Jing Chen (Nanfang Hospital, Southern Medical University) for providing clinical strains.

## DATA AVAILABILITY

The mass spectrometry proteomics data have been deposited to the ProteomeXchange Consortium via the PRIDE(Vizcaino et al., 2016) partner repository with the dataset identifier PXD030283 (Username, Password: l0o93cZD).

## AUTHOR CONTRIBUTIONS

X.S., R.G and Q.-Y. H., designed the project and revised the paper. Z.F., F.L., K.C., Z.Z., Z.F., L.C., and S.L.and Y.D. performed the experiment and data-analysis. Z.F., F.L. and K.C., wrote the manuscript.

---

**Figure 3—figure supplement 1.**
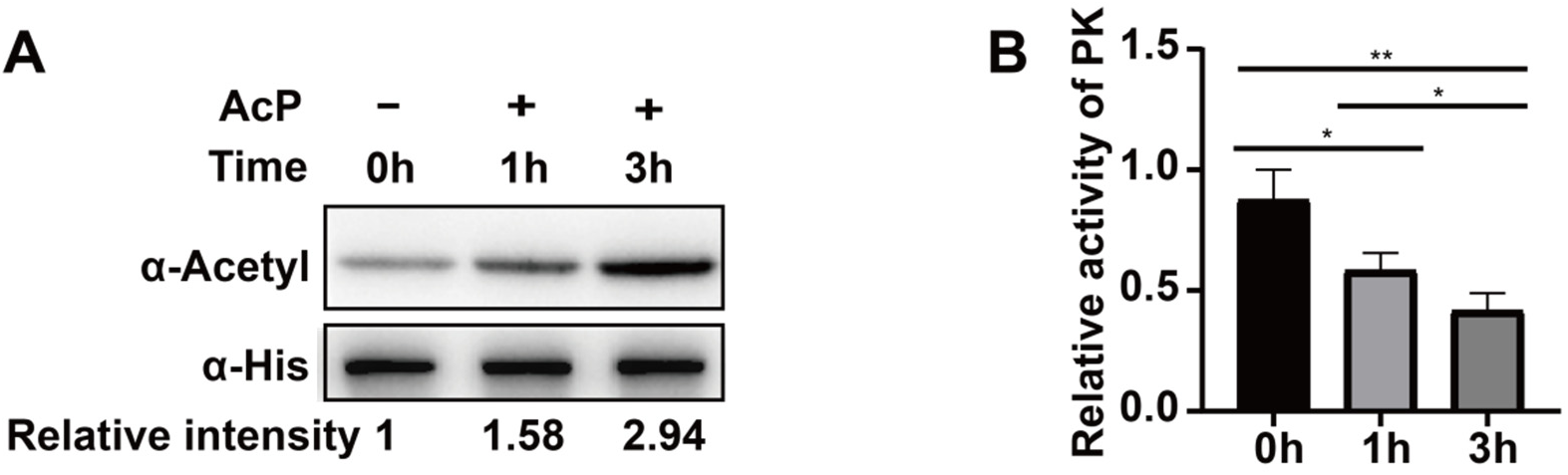
AcP acetylates PykF and reduces its enzyme activity. (A) AcP acetylates PykF in vitro. Western blots are representative of at least three independent replicates. (B) PykF enzyme activity detection.(n = 3 independent biological replicates). Two-tailed unpaired t-test was used for statistical analysis. (ns, no significant; *, p < 0.05; **, p < 0.01; ***, p < 0.005; ****, p < 0.001). Figure 3—figure supplement 1 source data 1. Uncropped blots for figure supplement 1A. Figure 3—figure supplement 1 source data 2. PykF enzyme activity for figure supplement 1B.

**Figure 5—figure supplement 1.**
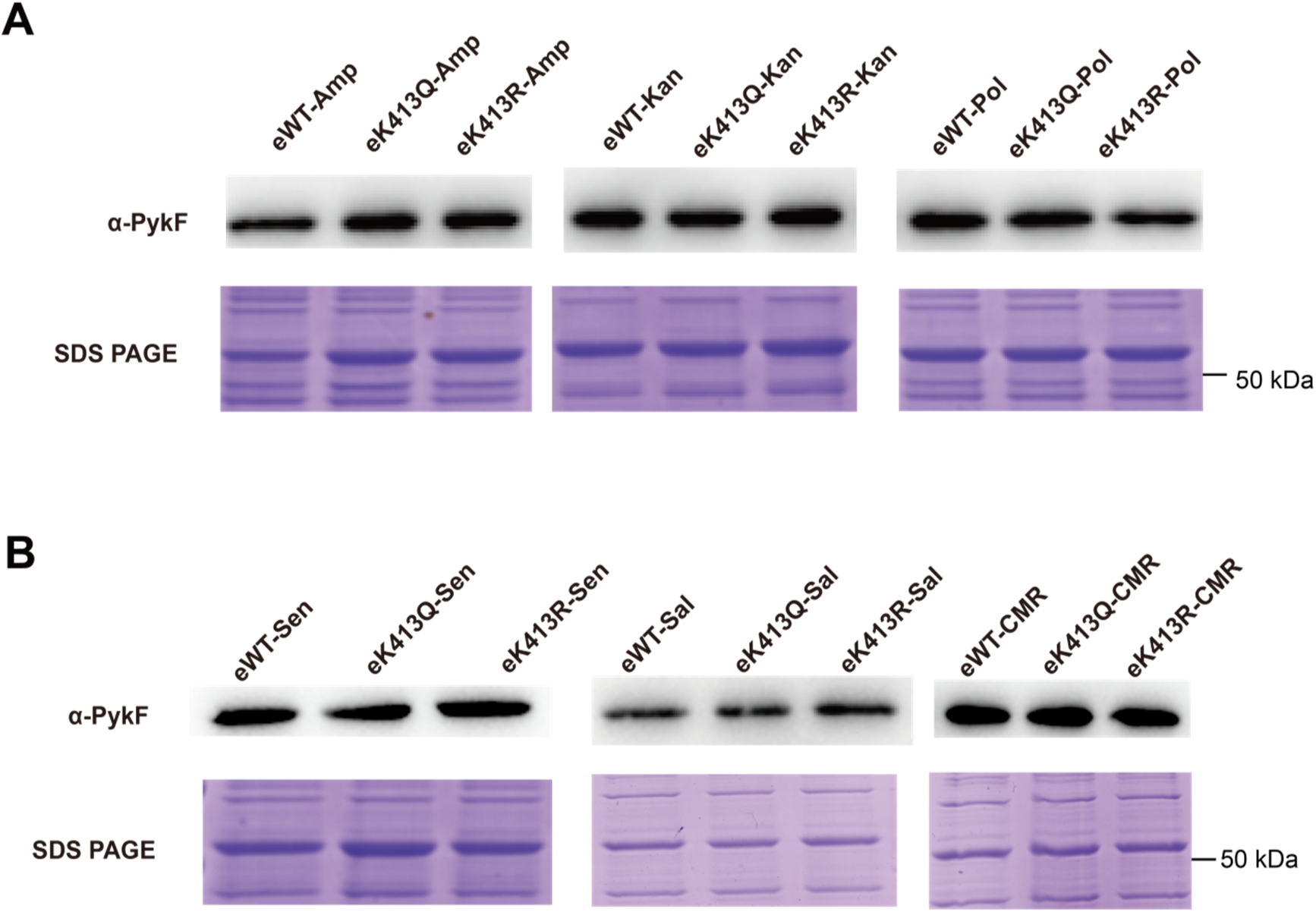
The stability of PykF expression in the complementary strains of E. coli (A) and *Salmonella* (B) were detected by western blot. Experiments were performed in triplicate. Figure 5—figure supplement 1 source data 1. Uncropped gels and blots for figure supplement 1.

